# Determinants of receptor tyrosine phosphatase homophilic adhesion: structural comparison of PTPRK and PTPRM extracellular domains

**DOI:** 10.1101/2022.06.23.497309

**Authors:** Iain M. Hay, Maria Shamin, Eve R. Caroe, Ahmed S. A. Mohammed, Dmitri I. Svergun, Cy M. Jeffries, Stephen C. Graham, Hayley J. Sharpe, Janet E. Deane

**Affiliations:** Cambridge Institute for Medical Research, University of Cambridge, Cambridge, CB2 0XY, UK; Signalling Programme, Babraham Institute, Babraham Research Campus, Cambridge CB22 3AT, U.K; MRC Laboratory of Molecular Biology, Cambridge CB2 0QH, U.K; Retroviral Replication Laboratory, Francis Crick Institute, London NW1 1AT, U.K; European Molecular Biology Laboratory (EMBL) Hamburg Site, Hamburg, Germany; Department of Pathology, University of Cambridge, Tennis Court Road, Cambridge CB2 1QP, UK

**Keywords:** cell adhesion, tyrosine‐protein phosphatase (tyrosine phosphatase), cell contact, receptor structure‐function, X‐ray crystallography, small angle X-ray scattering (SAXS)

## Abstract

The type IIB receptor protein tyrosine phosphatases (R2B RPTPs) are cell surface transmembrane proteins that engage in cell adhesion via their extracellular domains (ECDs) and cell signaling via their cytoplasmic phosphatase domains. The ECDs of R2B RPTPs form stable, homophilic, trans interactions between adjacent cell membranes. Previous work has demonstrated how one family member, PTPRM, forms homodimers; however, the determinants of homophilic specificity remain unknown. We have solved the X-ray crystal structure of the membrane-distal, N-terminal domains of PTPRK that form a head-to-tail dimer consistent with intermembrane adhesion. Comparison with the PTPRM structure demonstrates inter-domain conformational differences that may define homophilic specificity. Using small-angle X-ray scattering we determined the solution structures of the full-length ECDs of PTPRM and PTPRK, identifying that both are rigid, extended molecules that differ in their overall long-range conformation. Furthermore, we identify one residue, W351, within the interaction interface that differs between PTPRM and PTPRK and show that mutation to glycine, the equivalent residue in PTPRM, abolishes PTPRK dimer formation *in vitro*. This comparison of two members of the receptor tyrosine phosphatase family suggest that homophilic specificity is driven by a combination of shape complementarity and specific but limited sequence differences.

**SIGNIFICANCE STATEMENT:** Cell-cell contacts are dynamically regulated, in part, by the actions of tyrosine kinases and phosphatases. The R2B family of receptor tyrosine phosphatases combine an adhesive extracellular domain with intracellular catalytic domains that bind and dephosphorylate key cell adhesion and polarity proteins. Previous work demonstrated that the extracellular domains form head-to-tail homodimers but, as the interface was composed of residues conserved across the family, homophilic specificity determinants remained unclear. We have used a range of structural techniques including X-ray crystallography, small angle X-ray scattering and AlphaFold modelling to demonstrate that, despite their similarity, two members of the R2B family possess significant differences in their overall shape. Our results support that a combination of subtle shape and sequence variations may determine homophilic binding.

## INTRODUCTION

Cell-cell adhesion confers mechanical integrity to tissues that is critical for proper development and barrier function (1). Cell surface adhesion molecules hetero- or homodimerize to connect cells and are linked to the cytoskeleton via intracellular adhesive plaques. These adhesion complexes are highly dynamic and regulated by post translational modifications, in particular tyrosine phosphorylation (2). The actions of protein tyrosine kinases and phosphatases determine phosphotyrosine levels, which can be coupled to cell surface receptor domains, enabling responses to external signals and subsequent regulation of cell adhesion proteins (3,4). The type IIB receptor protein tyrosine phosphatases (R2B RPTPs) combine extracellular adhesion domains with intracellular catalytic phosphatase domains and may act as cell contact sensors in phosphorylation-based signaling events. R2B RPTPs function at cell contact sites and signal through the recruitment and regulation of multiple adhesion plaque protein substrates (5,6).

The four human R2B RPTPs (PTPRK, PTPRM, PTPRT and PTPRU) are type one transmembrane proteins and share a common extracellular domain (ECD) architecture of one MAM (meprin/A5/μ), one immunoglobulin (Ig)-like and four fibronectin type-III (FN) domains, with the most membrane proximal FN domain undergoing furin cleavage in the secretory pathway (7). This is followed by a transmembrane helix, an uncharacterized juxtamembrane domain and two tandem intracellular phosphatase domains (8). Previous studies have shown that PTPRK, PTPRM and PTPRT form homodimers, but not heterodimers, in cell aggregation assays (9–12). The homophilic (trans) interactions of R2B RPTPs determine their subcellular localization (5,11) and have been proposed to function as spacer clamps between cell membranes (13). Structural and biophysical studies demonstrate that for PTPRM the minimal unit required for dimerization in solution consists of the N-terminal MAM, Ig and first FN domain (hereafter referred to as MIFN1) (14). Dimer formation is pH-dependent, with the ECDs being monomeric at pH 6 and dimeric at pH 8 (14). This is thought to prevent formation of homophilic dimers within the secretory pathway, limiting dimerization to the cell surface at physiological pH (13).

Importantly, although the PTPRM ECD structure shed light on the nature of dimer formation, it did not explain the homophilic binding specificity of R2B receptors. Residues that were identified in the interface and demonstrated as necessary for dimer formation (including Y297, R239, R240 and R409) are conserved across the family (13). An earlier study, using chimeric ECD constructs and clustering assays demonstrated that the MAM domain was necessary but not sufficient for homophilic specificity (15). We therefore set out to determine ECD structures of another family member, PTPRK, to allow a detailed comparison of these dimeric complexes and determine which properties are driving binding specificity.

## RESULTS

To investigate dimerization of PTPRK *in vitro*, we expressed and purified both full-length PTPRK-ECD and a construct encoding only the N-terminal MAM-Ig-FN1 domains (PTPRK-MIFN1). One possibility for how homophilic interactions may be maintained is that each family member possesses a unique combination of predicted N-linked glycosylation sites (**Fig. S1**). Previous structural and functional studies with PTPRM used protein that had been deglycosylated or expressed in insect cell-based systems, resulting in non-native glycosylation (13,14). In order to test whether differential glycosylation may play a role in homophilic specificity we expressed PTPRK ECD constructs in the human HEK293-F cell line, with no subsequent deglycosylation, to maintain native glycans. To accurately determine the oligomeric state of these constructs and to test the pH sensitivity of dimerization, we performed size-exclusion chromatography coupled to multi-angle light scattering (SEC-MALS). Based on previous studies with PTPRM, we carried out SEC-MALS analysis at pH 8, where PTPRK would be expected to be dimeric, and pH 6 where monomers were anticipated. For each experiment, protein conjugate analysis was performed to calculate the proportional mass contribution of protein and glycan components. As expected, the full-length PTPRK-ECD was a pH sensitive homodimer, with total protein masses corresponding to dimer and monomer at pH 8 and pH 6, respectively (**Fig. 1A and B**). Similarly, the truncated N-terminal construct PTPRK-MIFN1 showed the same pattern of dimerization (**Fig. 1B and C**), confirming this fragment to be sufficient for formation of the pH sensitive homodimer.

**Figure 1.**
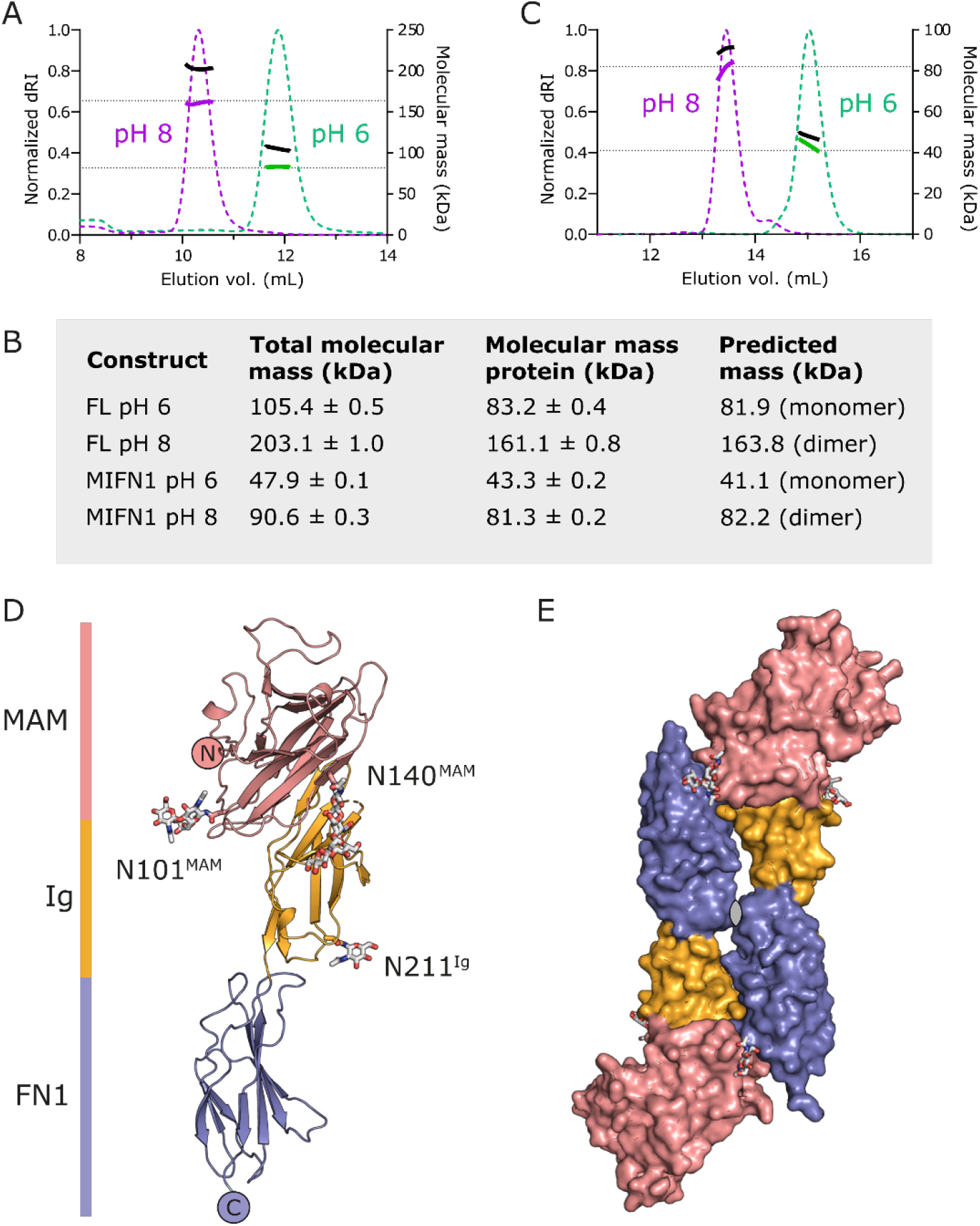
Structure of the PTPRK extracellular domain minimal dimerization unit. **(A)** SEC-MALS analysis of the full-length PTPRK-ECD. The SEC elution profile (normalized differential refractive index, dRI, dashed colored lines) and molecular mass distribution (total mass; black solid line, protein mass; colored solid line) at pH 8 (magenta) and pH 6 (green) are shown. Dashed horizontal lines indicated the predicted protein mass of PTPRK-ECD monomer and dimer. **(B)** Summary table of SEC-MALS mass determinations with and without conjugate analysis for attached glycans. **(C)** SEC-MALS analysis of the PTPRK-MIFN1 fragment, illustrated as described for A. **(D)** Ribbon diagram of the PTPRK-MIFN1 monomer, colored by domain. MAM: pink, Ig: orange, FN1: blue. N-linked glycosylation sites are labelled and glycans shown in stick representation. **(E)** Surface representation of the PTPRK-MIFN1 dimer, colored as in C. The 2-fold non-crystallographic symmetry axis is marked.

The PTPRK-MIFN1 fragment was crystallized, the X-ray structure solved by molecular replacement using the corresponding fragment of the PTPRM-ECD (PDB ID: 2V5Y, (13)), and the structure refined to 3.0 Å resolution (**Fig. 1D** and **Table S1**). The structure comprises a MAM domain (residues 31-192) that forms an extensive intramolecular interface with the Ig domain (residues 193-290), followed by the FN1 domain (residues 291-388) that makes few inter-domain contacts with the Ig domain (discussed below). Three glycosylation sites were observed in the electron density maps at asparagine residues N101 and N140 in the MAM domain and N211 in the Ig domain. A surface loop on the Ig domain encompassing residues 221-225, which possesses low conservation across the R2B family (**Fig. S1**), was unable to be built due to poor quality electron density. Two chains of PTPRK-MIFN1 were present in the asymmetric unit of the crystal structure forming a head-to-tail dimer and burying a surface area of ~1750 Å^2^(**Fig. 1E**). The dimer interface is formed between the MAM-Ig domain from one molecule and the FN1 domain of the opposing molecule, similar to that observed previously for PTPRM (13). Of the three observed glycosylation sites in PTPRK-MIFN1, N140 and N211 are located on the surface distal from the dimer interface and therefore are not likely to be important for determining homophilic binding specificity (Fig. 1D and E). N101 is located at the edge of the interface and may play a role in stabilization of the interaction.

To probe what might be contributing to binding specificity, the dimer interfaces were analyzed in further detail. The PTPRK-MIFN1 dimer is stabilized by several hydrogen bonds and salt bridges present at the MAM-Ig:FN1 interface (**Fig. 2A**). To aid understanding of these interactions the residue number will be accompanied by the domain they lie within, for example N207 in the Ig domain will be N207^Ig^. A loop encompassing residues 348-354 at the C-terminal end of the FN1 domain forms several key interactions. The side chain of K349^FN1^ forms hydrogen bonds with the opposing N101^MAM^, as well as the W160^MAM^ backbone carbonyl oxygen (**Fig. 2B**). Notably, the orientation of the W160^MAM^ carbonyl oxygen is caused by a *cis*-peptide bond with the adjacent P161^MAM^, a conformation that is stabilized by the side chain of W160^MAM^ stacking with the N-acetylglucosamine of the N101^MAM^ glycan. W351^FN1^ and H352^FN1^ both occupy a deep groove located at the MAM-Ig boundary of the opposing molecule, burying ~90% of their solvent accessible surface area. W351^FN1^ lies in a hydrophobic pocket created by the aliphatic portion of the R200^Ig^ side chain and F159^MAM^. R200^Ig^ additionally forms a hydrogen bond with the backbone carbonyl oxygen of K349^FN1^. H352^FN1^ occupies a hydrophilic pocket, forming hydrogen bonds with the side chains of T103^MAM^ and S157^MAM^, and the H352^FN1^ side chain imidazole ring stacks against the side chain of H197^MAM^. Finally, D354^FN1^ forms a salt bridge with R250^Ig^(**Fig. 2B**).

**Figure 2.**
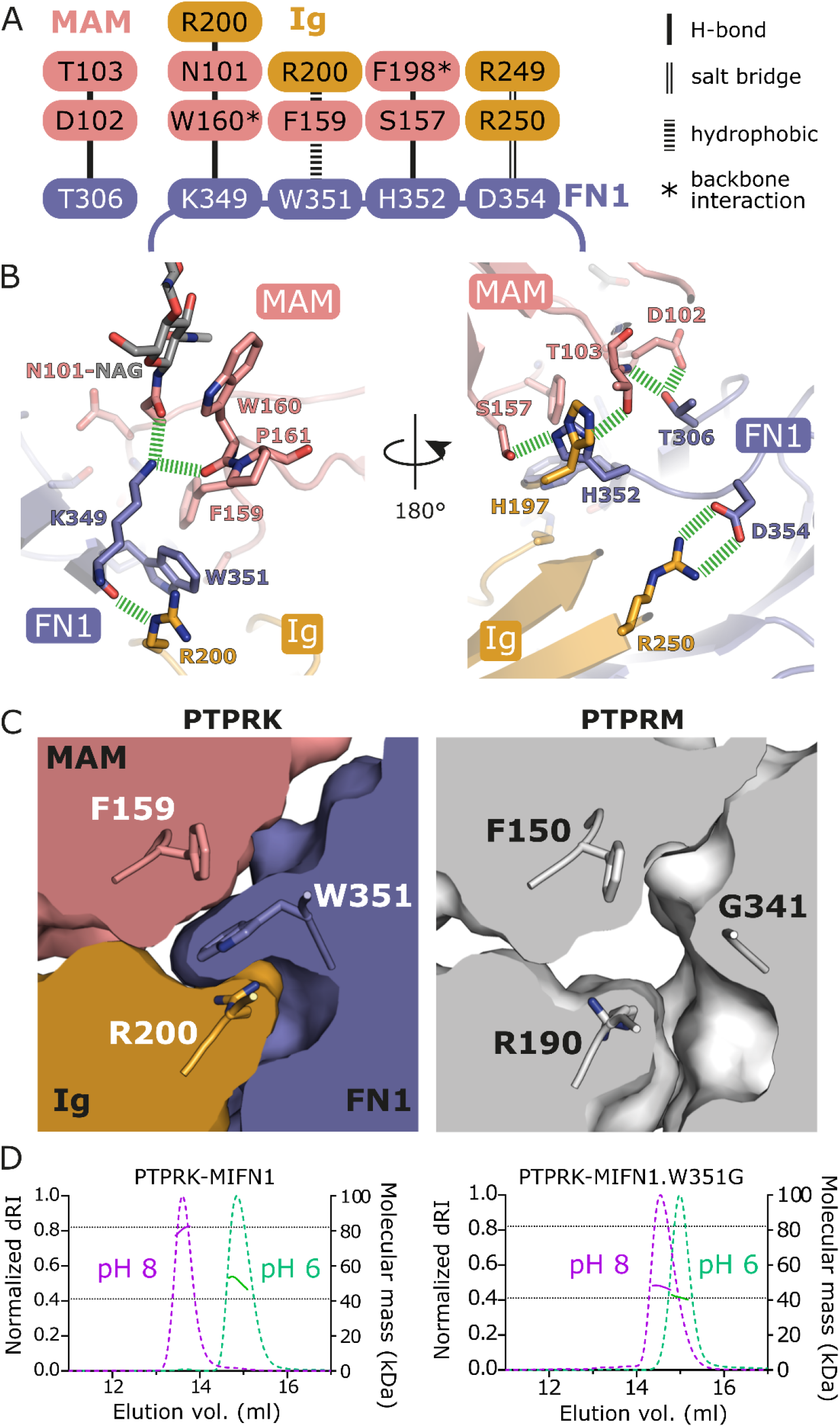
Dimerization interface of the PTPRK-MIFN1 structure. **(A)** Schematic diagram showing the hydrogen bonds, salt bridges and hydrophobic interactions at the MAM-Ig:FN1 interface within the PTPRK-MIFN1 homodimer. Stacked residues on the MAM-Ig side denote residues which have a common interacting residue in the FN1 domain. *denotes interactions formed by peptide backbone carbonyl oxygens. **(B)** Molecular detail of the PTPRK-MIFN1 homodimer interface. Side chains and backbone atoms of key interaction residues from (A) are highlighted, with hydrogen bonds and salt bridges highlighted (dashed green lines). The N-acetylglucosamine of the N101 linked glycan chain is shown (gray sticks). **(C)** Left: Cross-section of the MAM-Ig:FN1 interface in the PTPRK-MIFN1 dimer. W351 in the FN1 domain forms hydrophobic interactions with F159 and the aliphatic portion of R200 in the MAM and Ig domains, respectively. Right: Cross-section of the MAM-Ig:FN1 interface in the PTPRM-ECD dimer (PDB ID: 2V5Y) shown in the same orientation. **(D)** Left: SEC-MALS analysis of WT PTPRK-MIFN1. The SEC elution profile (normalized differential refractive index, dRI, dashed colored lines) and calculated protein mass distributions following conjugate analysis (colored solid line) are shown at pH 8 (magenta) and pH 6 (green). Dashed horizontal lines indicated the predicted protein mass of PTPRK-MIFN1 monomer and dimer. Right: SEC-MALS analysis of PTPRK-MIFN1.W351G, presented as for WT PTPRK-MIFN1 (left panel).

All but one of these key residues are completely conserved across the R2B receptor ECDs, the difference being that PTPRK W351^FN1^ is a glycine in PTPRM (**Fig. S1**). Despite lacking a bulky side chain at the equivalent position of W351, the binding groove between F159^MAM^ and R200^Ig^ that is occupied by this residue is maintained in PTPRM (**Fig. 2C**). To investigate the role of residue W351^FN1^ in dimer formation, we expressed and purified a W315G mutant form of PTPRK-MIFN1 for SEC-MALS analysis to test pH-dependent homodimer formation. Mutation of W351 in PTPRK to glycine was sufficient to inhibit dimerization, remaining primarily monomeric at pH 8 (**Fig. 2D**). Unfortunately, the reciprocal mutation G341W in the PTPRM-MIFN1 resulted in protein that aggregated to non-discrete oligomers in solution and was not suitable for further analysis. While these experiments demonstrate that W351 is required for PTPRK dimerization *in vitro*, they cannot explain the basis of homotypic selectivity as PTPRM has a surface cleft that could accommodate the W315 side chain. Furthermore, the other members of the R2B family, PTPRT and PTPRU both conserve this Trp residue suggesting that alternative structural differences may be driving homophilic specificity.

Detailed comparison of the PTPRK-MIFN1 structure with the equivalent region of PTPRM reveals a significant hinge movement (approximately 15°) of the FN1 domain relative to the MAM-Ig domains (**Fig. 3A**). This hinge movement is sufficient to induce a steric clash in the dimer interface when the MAM-Ig domains of PTPRK and PTPRM are superposed (**Fig. 3B**). This inter-domain movement may represent inherent flexibility between these domains or may represent rigid structural differences that contribute to homophilic specificity. For elongated multi-domain proteins, such as the R2B family, even small changes in inter-domain orientation have the potential to result in large conformational differences over the length of the full ECD. Superposing the PTPRK-MIFN1 structure on the PTPRM-ECD structure (encompassing the MAM-Ig-FN1-3 domains, PDB ID: 2V5Y), using the MAM-Ig domains for alignment, illustrates these potential differences (**Fig. 3C**). Models of the full-length ECDs of PTPRM and PTPRK generated using AlphaFold2 (AF2) possess significantly different inter-domain orientations relative to each other and also to the available crystal structures (**Fig. S2**). This lack of confidence in the inter-domain positions is quantified in the predicted alignment error (PAE) plots for these AF2 models (**Fig. S2B**). Despite this conformational range, the hinge movement observed in the PTPRK-MIFN1 structure determined here is not captured by the AF2 PTPRK models, suggesting the full-length ECDs may be even more conformationally distinct than is predicted by these models (**Fig. S2C**). These observations support the need for experimental approaches to determine the long-range shape of the ECD domains.

**Figure 3.**
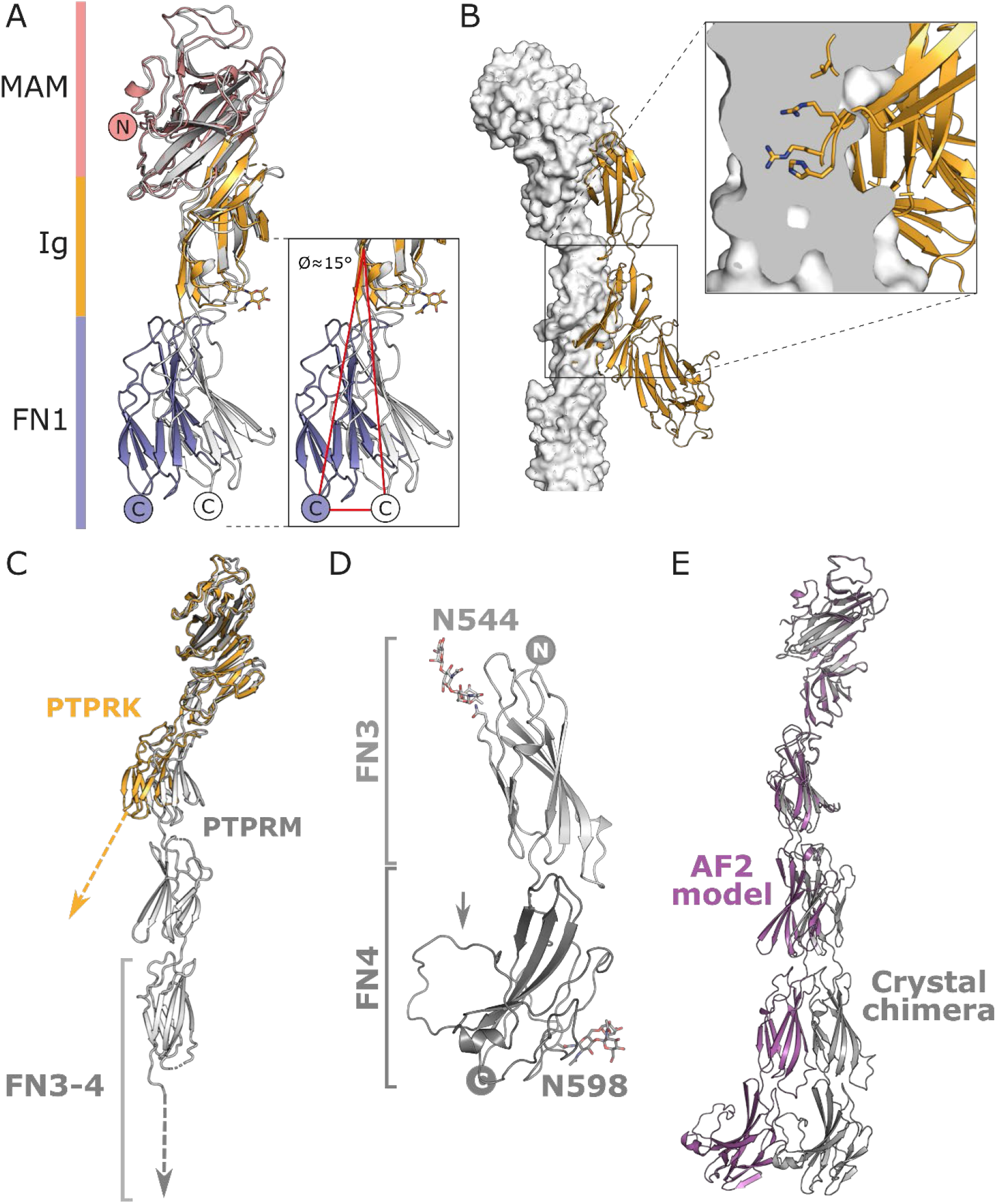
The PTPRK and PTPRM ECD structures possess different inter-domain conformations. **(A)** Ribbon diagram of a PTPRK-MIFN1 monomer, colored by domain. MAM: pink, Ig: orange, FN1: blue. Shown is a structural alignment over the MAM-Ig domains of the equivalent MIFN1 fragment from the full-length PTPRM-ECD structure (gray, PDB ID: 2V5Y). Inset: The FN1 domain of PTPRK possesses a ~15° shift from a hinge point at the Ig-FN1 boundary when compared to PTPRM. The glycan on PTPRK N211, which is absent in PTPRM, is shown (orange sticks). **(B)** The full-length PTPRM-ECD (gray surface) was aligned to one chain of the PTPRK-MIFN1 dimer (orange cartoon). Inset: Cross-section of the PTPRM-ECD overlay highlighting the extensive steric clashes when aligning PTPRM to the PTPRK dimer. **(C)** Ribbon diagram of a PTPRK-MIFN1 monomer (orange) aligned to the MAM-Ig domains of the PTPRM-ECD (gray). Dashed arrows highlight the potential for long-range structural divergence as a result of differing hinge angles at the Ig-FN1 boundary. **(D)** Ribbon diagram of the PTPRM-FN3-4 structure. N-linked glycosylation sites are labelled and glycans shown in stick representation. The loop in FN4 containing the furin cleavage site is labelled (arrow).**(E)** Comparison of structures for the full-length ECD of PTPRM determined using available crystal structures (chimera, gray) and AlphaFold2 (AF2, purple). Structures were overlaid using the MAM-Ig domains only.

The available X-ray structure of the PTPRM ECD (PDB ID: 2V5Y, (13)) is missing the final FN4 domain, limiting our understanding of the relative orientation of this domain. We therefore determined the crystal structure of the PTPRM-FN3-4 domains (**Fig. 3D** and **Table S1**). Crystals grew in two space groups, *P*2_1_2_1_2_1_ and *P*3_2_21, possessing two and four molecules per asymmetric unit, respectively (**Fig. S3A and B**). All six chains are essentially identical in structure (pairwise RMSDs over 220 Cα atoms range from 0.44 to 0.92 Å, **Fig. S3C**). The main difference between the protein structures is that one chain in the *P*2_1_2_1_2_1_ crystal has a disordered loop (residues 628–641) that in all other chains was ordered via intermolecular interactions (**Fig. S3A and B**). This loop is anchored at one end via a disulfide bond within the FN4 domain (C642 to C716) and the sequence encompasses the furin-cleavage site of PTPRM, illustrating that this site is surface accessible within a conformationally dynamic loop but that cleavage would not significantly destabilize the fold of the FN4 domain (**Fig. 2D**). There are two predicted glycosylation sites in FN3−4, one in each domain: N544 in FN3; and N598 in FN4. In both crystal forms one chain is missing the glycan on N598, indicating that PTPRM-FN4 not only adopts conformational differences in the furin-cleavage loop but can also undergo heterogeneous glycosylation.

Although the previous structure of the PTPRM ECD had no interpretable electron density for the FN4 domain, it was present in the construct. Superposition of the FN3 domains from our structure with this previous PTPRM structure (13) demonstrates that the FN4 domain can be accommodated without significant steric clashes with surrounding chains, and FN4 co-localizes with weak and disconnected electron density that is unmodeled in the deposited structure. This is consistent with the FN3−4 domains adopting similar conformations when crystallized in isolation or as part of the full-length ECD. This alignment therefore allowed the generation of a full-length ECD chimeric structure based on crystallographic data (**Fig. 3E**). Superposition of this chimeric PTPRM structure and the full-length PTPRM-ECD AF2 models (aligned using the MAM-Ig domains) reveals the extent to which these models for the exact same protein can diverge (**Fig. 3E**). This highlights the difficulty in confidently determining long-distance conformational differences between PTPRM and PTPRK ECDs from crystallographic snapshots or *ab initio* models alone.

To probe the overall shape of these domains in solution we implemented small-angle X-ray scattering coupled to SEC and MALS (SEC-SAXS-MALS). The structural parameters determined for purified PTPRK and PTPRM ECDs confirm that both proteins elute as glycosylated monomers at pH 6 (**Table S2**). The experimental SAXS profiles for PTPRK-ECD (**Fig. 4A**) and PTPRM-ECD (**Fig. 4B**) appear similar, indicating conserved global structural features in both proteins. Each of the ECDs are extended particles and both display highly skewed real-space distance distributions (*p*(*r*) profiles) with a maximum particle dimension, *D_max_*, of 26 nm (**Fig. 4C**). Importantly, the *p*(*r*) profiles at shorter distances are not identical, confirming differences in the local domain organization of these proteins. An overlay of the scaled SAXS data (**Fig. S4A**) also shows deviations between the PTPRK and PTPRM scattering intensities at lower angles, in particular spanning an *s*-range of 0.5–1.6 nm^−1^, which corresponds to real-space distances of 3.9–12.5 nm. These observations, combined with a consistent 0.2 nm increase in the radius of gyration (*R_g_*) and 0.1 nm decrease in the *R_g_* of cross-section (*R_g_*^c^) of PTPRM compared to PTPRK (**Table S2; Fig. S4B**), reveal distinct differences between the spatial disposition of the domains in the two constructs. With respect to the overall structural sampling of the ECDs, the dimensionless Kratky plots of the SAXS data (**Fig. S4C**) display a monotonic increase for *sR_g_* < 5, typical of particles with extended but relatively rigid conformations and not of proteins with domains connected by flexible linkers sampling diverse states in solution (16). Therefore, the differences seen in the *p*(*r*) profiles between PTPRM and PTPRK ECDs are not due to flexibility but driven by differences in the shape of these domains.

**Figure 4.**
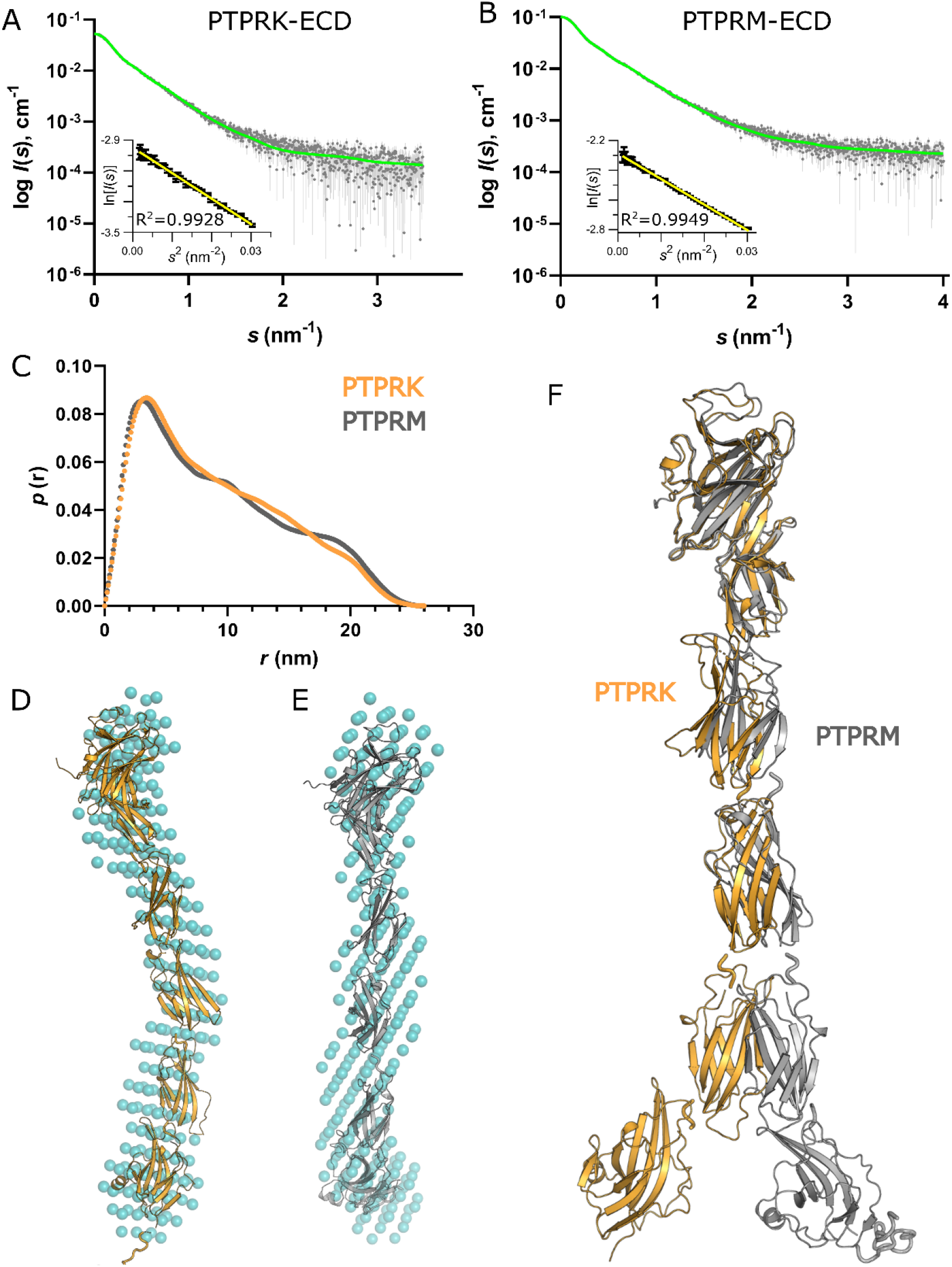
Small angle X-ray scattering data and ECD modelling. Averaged SEC-SAXS profile of **(A)** PTPRK-ECD and **(B)** PTPRM-ECD (gray diamonds) and corresponding fit against the data of respective pseudo-atomic models shown in (**D**) and (**E**) (green line). *Inset*, Guinier plot of ln*I* (*s*) versus *s*^2^ for *sR_g_* < 1.3. **(C)***p*(*r*) versus *r* profiles of PTPRK-ECD (orange) and PTPRM-ECD (gray)**. (D)** The *ab initio* dummy-atom bead model of PTPRK-ECD calculated using DAMMIN (cyan spheres) overlaid with the best pseudo-atomic model (lowest χ^2^) generated by fitting to the SAXS data. **(E)** As for panel D but for PTPRM-ECD data. **(F)** A spatial comparison of the PTPRK-ECD (orange) and PTPRM-ECD (gray) SAXS-based models aligned relative to the N-terminal MAM-Ig domains.

A*b initio* shape analysis with the program DAMMIN (17) yielded narrow (~3–4 nm), extended and slightly bent dummy atom models for both ECDs (**Fig. 4D and E**). Pseudo-atomic models of the PTPRK and PTPRM ECDs were generated by combining the available crystal structures with AF2 predictions for uncharacterized domains, with glycans added to the relevant residues. Initial hybrid models for the PTPRK and PTPRM ECDs fit poorly to the SAXS data, with discrepancy χ^2^ = 1.78 and probability of systematic deviations between the model fit and the scattering data (CorMap *P,* (18)) of 5.0×10^−6^ for PTPRK and χ^2^ = 1.42 (CorMap *P* = 1.0×10^−8^) for PTPRM. Rigid body refinement of these initial models to the SAXS data using CORAL (19), which allowed repositioning of the MAM-Ig and FN domains with respect to each other, yielded pseudo-atomic models with significantly improved fits, χ^2^ = 1.17 (CorMap *P* = 0.072) for PTPRK-ECD and χ^2^ = 1.10 (CorMap *P* = 0.293) for PTPRM-ECD (**Fig. 4D and E, Table S2**). Superposition of these final pseudo-atomic models (**Fig. 4F**) confirms that both adopt extended conformations in solution, but that PTPRK appears more ‘bent/twisted’ compared to PTPRM. Such conformational differences are consistent with the observed differences in the *p*(*r*) profiles for PTPRK and PTPRM, and with the smaller radius of gyration (*R_g_*) and larger *R_g_* cross-section (*R*_g_^c^) obtained from the SAXS data for PTPRK. Furthermore, the conformations of the PTPRM and PTPRK ECDs differ in shape sufficiently that the SAXS-based hybrid model of one construct does not fit the SAXS data for the other (PTPRK-ECD, *χ*^2^PTPRM-SAXS = 1.75, CorMap *P* = 9.8×10^−12^; PTPRM-ECD, *χ*^2^PTPRK-SAXS = 1.26, CorMap *P* = 3.6×10^−5^). These SAXS models indicate that the overall shape of these R2B ECDs differ from each other, which may drive homophilic specificity.

To probe what might be contributing to the different shapes of PTPRK and PTPRM ECDs, the X-ray structures were analyzed in closer detail with particular focus on the intramolecular contacts between the MAM-Ig and FN1 domains. Comparative analysis of PTPRK and PTPRM shows a large degree of conservation at the intramolecular interface between the Ig and FN1 domains (**Fig. S5**). In PTPRK, the side chain of N207^Ig^ forms hydrogen bonds with the side chain of R367^FN1^ and the backbone carbonyl oxygens of P368^FN1^ and E370^FN1^. The former two bonds are conserved in PTPRM and inspection of electron density maps from PDBe (20,21) supports the third also being conserved. The PTPRK D320^FN1^ side chain and backbone amide nitrogen form hydrogen bonds with the backbone amide nitrogen and carbonyl oxygen, respectively, of E290^Ig^. Inspection of PTPRM electron density maps also supports this interaction. One subtle difference is that in PTPRM E195^Ig^ forms a salt bridge with R357^FN1^, whereas in PTPRK the equivalent E205^Ig^ residue instead forms an intradomain salt bridge with R289^FN1^, which is a lysine residue in PTPRM. Furthermore, on a loop of the PTPRK Ig domain, near the FN1 interface, there is an N-linked glycan at N211 that is absent in PTPRM (**Fig. 3A**). This glycan sits near the apex of the hinge movement of FN1 and lies on the opposite side of PTPRK from the dimer interface. Although the modelled glycan does not make any stable contacts with the FN1 domain, the presence of a bulky glycan tree here could cause the FN1 domain to ‘swing’ away from the Ig domain. Similar steric hindrance based conformation changes have been observed previously (22,23), for example, in the glycan-mediated regulation of plasminogen activator inhibitor−1 (24).

## DISCUSSION

The ECDs of R2B RPTPs engage in homophilic (trans) interactions to bridge plasma membrane contact sites. Previous studies using PTPRM identified how these ECDs form head-to-tail dimers via the N-terminal MAM-Ig-FN1 domains but did not investigate how homophilic specificity is determined. Here we determined the X-ray structure of this region of a related family member, PTPRK, allowing direct comparison with the PTPRM structure. Analysis of the crystallographic PTPRK and PTPRM homodimers identified that the vast majority of residues involved in the dimerization interface are conserved across the R2B family of proteins, indicating that any contributions of primary sequence composition to determining homophilic binding specificity are likely to be subtle. Indeed, previous work identifying the sequence determinants of dimer formation using cell clustering assays mutated residues that are conserved across the R2B family, supporting that the crystallographic dimer is the relevant complex but not how homophilic specificity is established. The pH-dependency of dimer formation shown here for PTPRK is the same as that for PTPRM, supporting that both proteins possess similar electrostatic complementarity. Although a previous study suggested that surface charge may drive homophilic specificity (9), the electrostatic surfaces of PTPRM and PTPRK across the interaction interface appear very similar (**Fig. S6**). These observations suggest that the mechanism of homophilic specificity is not strongly driven by either sequence composition or surface charge differences.

Importantly, detailed analysis of the X-ray structures revealed differences in the relative orientations of the N-terminal MAM-Ig and FN1 domains that mediate dimerization. The interdomain contacts in PTPRK versus PTPRM suggest that the hinge movement between the MAM-Ig and FN domains is not driven by substantial sequence differences, but is instead driven by a small number of subtle differences that position the PTPRK FN1 domain such that it is likely to be structurally incompatible with forming a heterodimer with PTPRM. In support of this steric hindrance model, SAXS-based structural analysis of the full-length ECDs of PTPRK and PTPRM confirm that these proteins adopt rigid, extended but different conformations in solution. Fitting of crystallographic and AlphaFold models of PTPRK and PTPRM domains to the SAXS data reveal that the PTPRK ECD adopts a more bent/twisted shape when compared with the PTPRM ECD. These data support that, despite only subtle differences in their sequence composition, PTPRK and PTPRM ECDs adopt distinct overall shapes that are likely to favor homophilic, and hinder heterophilic, interactions.

Comparison of the X-ray and SAXS structures of the PTPRK and PTPRM ECDs support an important role for overall shape in homophilic specificity. There are two additional members of this family, PTPRT and PTPRU. As observed in this work, AF2 models for these large multi-domain ECDs cannot, on their own, provide direct insights into the mode of homophilic binding and cannot reliably predict overall shape. However, models of these proteins do allow for the analysis of surface properties such as electrostatics as done previously (9) and also repeated here using AF2 (**Fig. S6**). Interestingly, electrostatic surface representations for the ECDs of PTPRT and PTPRU do reveal some differences within the dimerisation interface. Future work including detailed structural analysis of these additional members of the R2B family will help shed more light on the exact nature of homophilic specificity in this family. Finally, the structural observations made here have been determined with isolated ECDs separate from their associated membrane domains. How these proteins are oriented on the cell surface is highly challenging to measure but may further contribute to homophilic specificity of R2B family members in the context of a membrane interface.

## EXPERIMENTAL PROCEDURES

### Plasmids and constructs

Amino acid numbering for DNA constructs is based on the following sequences; human PTPRK (Uniprot Q15262−3) and human PTPRM (Uniprot P28827−1). For recombinant expression in mammalian cells, PTPRK full ECD (residues 28−752) and MIFN1 (residues 28−385) and PTPRM full ECD (residues 19−742) and FN3−4 (residues 478−723) were PCR amplified and cloned into the pHLSec vector using AgeI and KpnI restriction sites allowing in-frame expression with an N-terminal secretion signal and a C-terminal Lys-His_6_tag.

### Protein expression and purification

Expression of recombinant extracellular proteins was performed by transient transfection in HEK−293F cells using polyethylenimine (PEI, Sigma Aldrich). 300 mL of culture at a density of 1.0 × 10^6^cells/mL was transfected with 300 μg DNA using 450 μg PEI. Culture medium was harvested 72 h post-transfection and secreted His-tagged proteins purified using Ni-NTA agarose (Qiagen) in batch mode. Beads were packed into a 20 mL gravity column, washed with 20 mL of wash buffer (50 mM Tris-HCl, pH 8, 500 mM NaCl, 20 mM imidazole) and eluted using wash buffer containing 250 mM imidazole. Eluted protein was further purified by size exclusion chromatography (SEC). For PTPRK and PTPRM ECD constructs SEC was performed using a HiLoad Superdex 200 pg 16/600 column (Cytiva) equilibrated in 100 mM Tris-HCl, pH 8, 250 mM NaCl. For the PTPRM FN3−4 constructs SEC was run on a HiLoad Superdex 75 pg 16/600 column (Cytiva) equilibrated in 20 mM Tris-HCl, pH 7.4, 150 mM NaCl.

### Multi-angle Light Scattering (MALS)

MALS experiments were performed immediately following SEC (SEC-MALS) by inline measurement of static light scattering (DAWN 8+; Wyatt Technology), differential refractive index (Optilab T-rEX; Wyatt Technology), and UV absorbance (1260 UV; Agilent Technologies). Samples (100 μl) at 1 mg/mL were injected on to a Superdex 200 Increase 10/300 GL column (Cytiva) equilibrated in pH 8 purification buffer (100 mM Tris-HCl, pH 8, 250 mM NaCl) at a flow rate of 0.4 mL/min. The molar masses of the major SEC elution peaks were calculated in ASTRA 6 (Wyatt Technology) using a protein dn/dc value of 0.185 mL/g. For determination of protein and glycan fractions, conjugate analysis was performed in ASTRA 6, using a glycan (modifier) dn/dc = 0.14 mL/g and theoretical UV extinction co-efficients calculated using ProtParam (25). For experiments at pH 6, ECD peak fractions were collected from pH 8 experiments and buffer exchanged using centrifugal concentration units (Amicon, Merck) into pH 6 purification buffer (50 mM MES, pH 6, 250 mM NaCl). SEC-MALS experiments at pH 6 were then carried out as for pH 8 experiments.

## Crystallization

Crystallization experiments were carried out in 96-well nanolitre-scale sitting drops (200 nL of purified protein with 200 nL of precipitant) equilibrated at 20°C against 80 μL reservoirs of precipitant. PTPRK-MIFN1 crystallization was performed using 9.5 mg/mL protein and diffraction quality crystals grew against a reservoir of 100 mM HEPES, pH 7, 10% (w/v) polyethylene glycol (PEG) 6000. Two different PTPRM FN3−4 crystals grew in related conditions. The *P*2_1_2_1_2_1_ crystals grew following equilibration of purified protein at 11.2 mg/mL against a reservoir containing 200 mM ammonium nitrate and 20% (w/v) PEG 3350. The *P*3_2_21 crystals were grown following microseeding as described previously (26) using a seed stock made from the *P*2_1_2_1_2_1_ crystals combined with purified protein at 9.1 mg/mL against a reservoir containing 100 mM ammonium nitrate and 10% PEG 3350. All crystals were cryoprotected in reservoir solution supplemented with 20% (v/v) glycerol and flash-cooled by plunging into liquid nitrogen.

### X-ray data collection and structure solution

X-ray diffraction datasets were recorded at Diamond Light Source (DLS) on beamlines I03, I04 and I04−1 (**Table S1**). Diffraction datasets were indexed and integrated using the XIA2 DIALS pipeline (27,28). For PTPRK-MIFN1, the initial structure was solved by molecular replacement (MR) using Phaser (29), with the MAM, Ig-like and first FN domains of the human PTPRM full ECD (PDB ID: 2V5Y, (13)) as a search model. For PTPRM-FN3−4, there was no model available (at the time) that encompassed both domains. Therefore, initial MR was carried out in Phaser searching for 2 copies of FN3 domain using residues 461−563 of the PDB-REDO model of 2V5Y. To model the FN4 domain, an alignment using HHPRED identified domains from 2M26, 3UTO and 1WFT that were used to generate an ensemble of models in Sculptor (30). This ensemble was used in Phaser, searching for 2 copies. The resulting solution was refined using autobuster (GlobalPhasing) and manually rebuilt using COOT. This was followed by density modification using Parrot (31) and NCS averaging of autobuster phases followed by autobuilding using Buccaneer (32). Further refinements of all structures were performed using COOT (21), ISOLDE (33) and phenix.refine (34). All graphical figures were rendered in PyMol (Schrödinger LLC) except for surface electrostatic images which were illustrated using ChimeraX (35). The atomic coordinates and structure factors have been deposited in the Protein Data Bank, www.pdb.org under accession codes 8A1F (PTPRK-MIFN1), 8A16 and 8A17 (PTPRM-FN3−4).

### AlphaFold2 Multimer structure predictions

All AF2 models were generated using default parameters and run via a locally installed version of AF2 (36). All models and associated statistics have been deposited in the University of Cambridge Data Repository (https://doi.org/10.17863/CAM.84929).

### Small-angle X-ray Scattering

SAXS experiments were performed using SEC-SAXS at the EMBL-P12 bioSAXS beam line (PETRAIII, DESY, Hamburg, Germany) (18) with inline MALS, refractive index and UV detectors (Wyatt miniDAWN® TREOS®, a Wyatt Optilab T-rEX (RI) refractometer and Agilent variable wavelength UV-Vis detector recording at 280 nm) (37). The SEC-SAXS-MALS data were recorded as detailed in **Table S2**. For data collection, 40 μL of PTPRK-ECD (2.3 mg/mL) and 30 μL of PTPRM-ECD (7.0 mg/mL) were injected at 0.35 mL/min onto an S200 Increase 5/150 column (Cytiva) equilibrated in 50 mM MES pH 6.0, 250 mM NaCl, 3% (v/v) glycerol. The SAXS data were recorded on a Pilatus 6M detector as a set of 2880 2D-data frames with 0.25 s exposure through the entire column elution. The 2D-to−1D azimuthal averaging was performed using the SASFLOW pipeline (18). The subtraction of appropriate buffer scattering intensities from the sample scattering collected through the single elution peak of either protein was performed using CHROMIXS (18,38). The processing and analysis of the final scaled and averaged SAXS data were performed using the ATSAS 3.0.2 software package (39). The extrapolated forward scattering intensity at zero angle, *I*(0), and the radius of gyration, *R_g_*, were calculated from the Guinier approximation (ln*I*(*s*) vs *s*^2^, for *sR_g_* < 1.3). The radius of gyration of cross section, *R_g_*^c^, was estimated using a modified Guinier plot (ln(*I*(*s*)s) vs *s*^2^, in the *s*^2^ range between 0.025–1.0 nm^−2^), while the dimensionless Kratky plots ((*sR_g_*)^2^*I*(*s*)/*I*(0) vs *sR_g_*) were generated as described in (16). Shape classification was performed using DATCLASS (40). The distance distributions in real-space *p*(*r*) were calculated using GNOM (41). A concentration-independent estimate of the molecular weight, extracted directly from the SAXS data, was determined using a Bayesian consensus method (42), while MALS-RI-UV conjugate molecular weight validation was performed as described above (41−43).

*Ab initio* modeling was performed using multiple individual DAMMIN shape reconstructions followed by spatial alignment and bead-occupancy/volume correction to generate both an averaged spatial representation and a final single dummy atom model for each protein (17,44). Atomistic models for the full ECDs of PTPRM and PTPRK were generated by combining available crystal structures with AF2 models for missing regions. Specifically, for PTPRM the available X-ray structures (PDB ID 2V5Y and the FN3−4 structure determined here) were combined following superposition of the FN3 domains (present in both models). Short stretches of missing residues (19−21 and 725−752) were added from the top-ranked AF2 model. For PTPRK, the MIFN1 X-ray structure determined here replaced the equivalent region in the top AF2 model. For both ECD models, the C-terminal His_6_ tag and hydrogens were added as well as complex mammalian glycans using the GLYCOSYLATION module of ATSAS in combination with the carbohydrate builder from the Glycam server (https://glycam.org/cb/). Subsequent rigid-body modelling of the domain orientations to the SAXS data was performed using CORAL (19). Five rigid bodies were defined: the MAM plus Ig domains together and the four FN domains independently. The final atomistic model fits to the SAXS profiles were calculated using CRYSOL (35 spherical harmonics, 256 points, with constant adjustment and the inclusion of explicit hydrogens) (45) and the quality evaluated using the reduced χ^2^ test and CorMap *P*-value. The SEC-SAXS and SEC-MALS data as well as the SAXS data modeling and analysis are made available in the Small Angle Scattering Biological Data Bank (SASBDB) (46), with the accession codes: SASDPF3 (PTPRK-ECD) and SASDPG3 (PTPRM-ECD) (19,45).

## Acknowledgements

We acknowledge Diamond Light Source for time on beamlines I03, I04 and I04−1 under proposal MX15916 and MX21426. Remote access was supported in part by the EU FP7 infrastructure grant BIOSTRUCT-X (Contract No. 283570). The synchrotron SAXS data was collected at beamline P12 operated by EMBL Hamburg at the PETRA III storage ring (DESY, Hamburg, Germany) with support from iNEXT-Discovery, project number 871037, funded by the Horizon 2020 programme of the European Commission. We thank Radu Aricescu for sharing DNA constructs of PTPRM. A Titan V graphics card used for this research was donated to S.C.G. by the NVIDIA Corporation. For the purpose of open access, the author has applied a Creative Commons Attribution (CC BY) licence to any Author Accepted Manuscript version arising from this submission.

## Funding Information

This study was supported by a Sir Henry Dale Fellowship jointly funded by the Wellcome Trust and the Royal Society awarded to H.J.S. (109407/Z/15/Z). J.E.D. was supported by a Royal Society University Research Fellowship (UF100371) and a Wellcome Trust Senior Research Fellowship (219447/Z/19/Z). I.M.H. was funded by a CIMR PhD studentship and Babraham Institutional bridging funds. E.R.C. was supported by a Royal Society Enhancement Award (RGF\EA\180151) and M.S. was supported by a Wellcome Trust PhD studentship (203984). A.S.A.M. was supported by a Deutsche Forschungsgemeinschaft grant (SV 9/11−1). H.J.S. is an EMBO Young Investigator and Lister Institute Prize Fellow.

## Data availability

The atomic coordinates and structure factors have been deposited in the Protein Data Bank, www.pdb.org under accession codes 8A1F (PTPRK-MIFN1), 8A16 and 8A17 (PTPRM-FN3−4). AF2 models and associated statistics have been deposited in the University of Cambridge Data Repository (https://doi.org/10.17863/CAM.84929). The SEC-SAXS-MALS data and models are available in the Small Angle Scattering Biological Data Bank (SASBDB) (46), with the accession codes: SASDPF3 (PTPRK-ECD) and SASDPG3 (PTPRM-ECD).

## Conflict of Interest

The authors declare they have no conflict of interest.

## Notes

https://doi.org/10.17863/CAM.84929

### Competing Interest Statement

The authors have declared no competing interest.

### Summary of Updates

Minor changes to title and abstract to clarify the comparisons are focussed on differences between PTPRK and PTPRM rather than all R2B family members. Addition of Supp Figure 6 showing models of the other family members PTPRU and PTPRT.

