## Supporting Information for "Determinants of receptor tyrosine phosphatase homophilic adhesion: structural comparison of PTPRK and PTPRM extracellular domains"

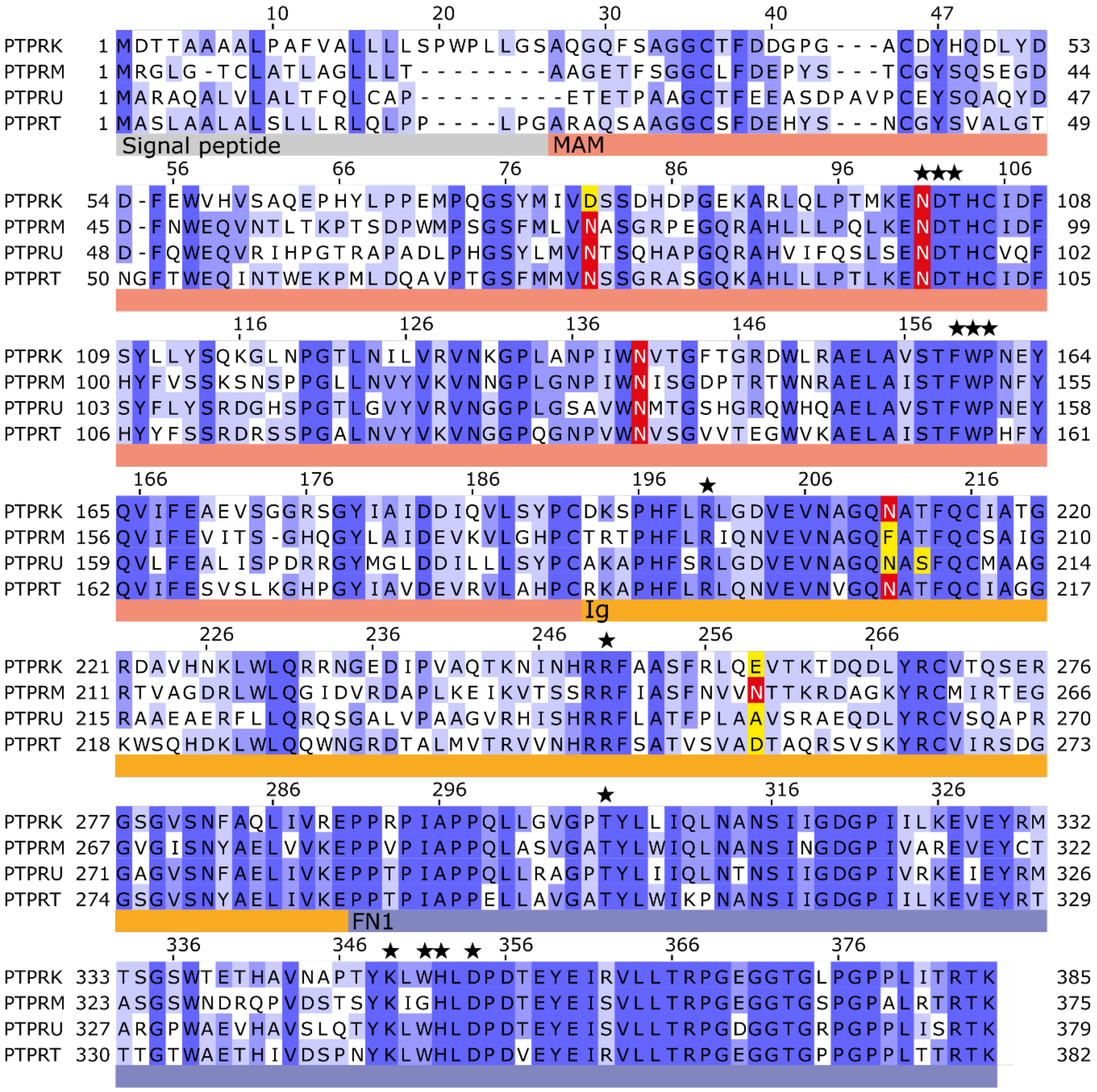


**Fig. S1. Sequence alignment of the MAM, Ig and FN1 domains of the ECDs from the four R2B PTP family members.** Multiple sequence alignment of the MAM (pink), Ig-like (Ig, orange) and first fibronectin type-III (FN1, blue) domains of the human R2B RPTPs, coloured by percentage identity (white-blue, 0-100% identity). Predicted N-glycosylation sites were identified using the NetNGlyC server (1) and are highlighted in red, with non-conserved sites in yellow. Residues involved in key intermolecular interactions at the MIFN1 dimer interface are highlighted (stars).

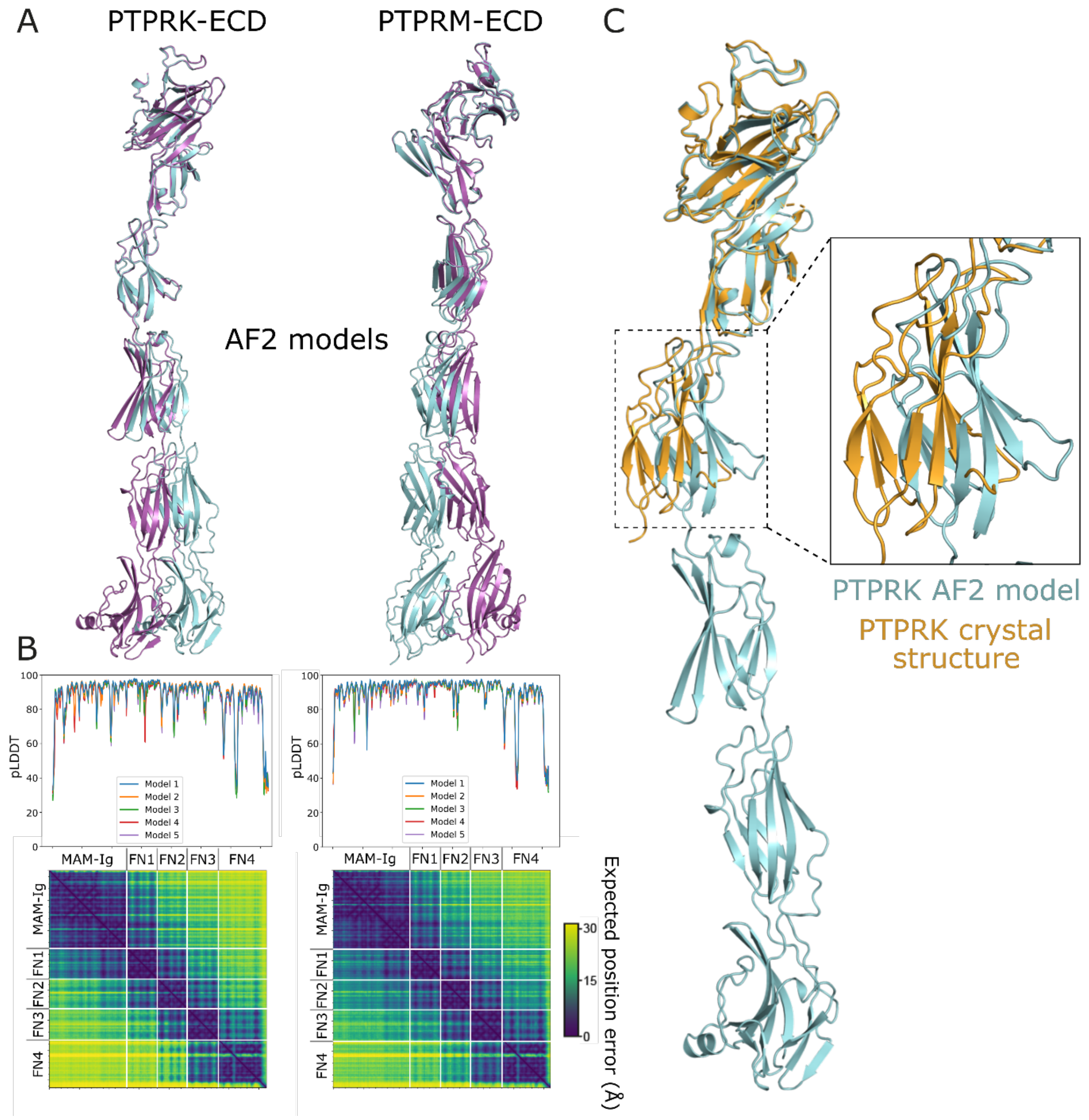


**Fig. S2. AlphaFold2 (AF2) models of the PTPRK and PTPRM ECDs. A.** For the PTPRK-ECD (left) and PTPRM-ECD (right) the top five AF2 models were superposed using the MAM-Ig domains as the reference for alignment. Two of the five models are displayed for clarity to demonstrate the range of long-distance conformations present in the ensemble of predicted structures (cyan and magenta). **B.** *Top,* pLDDT plots demonstrating good per-residue confidence for all PTPRK and PTPRM ECD models. *Bottom*, Representative Predicted Aligned Error (PAE) plots for PTPRK and PTPRM, with domain boundaries marked (white lines), demonstrating the low confidence (high error, yellow) of long-distance predictions. **C.** Alignment of the AF2 model for the PTPRK-ECD with the crystal structure of the PTPRK-MIFN1 using the MAM-Ig domains as the reference for alignment.

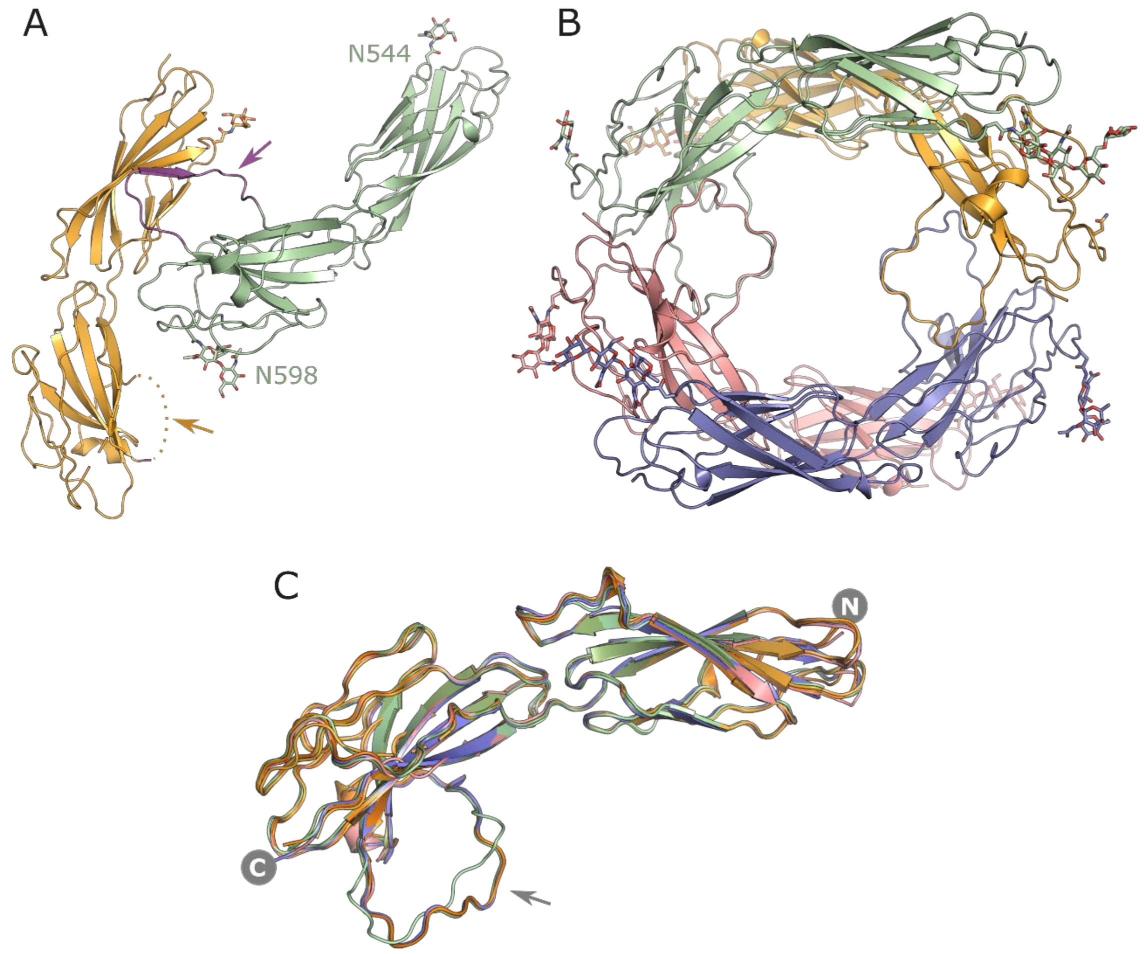


**Fig. S3. Tertiary assemblies of the PTPRM FN3-4 structures determined in two different spacegroups. A.** In $P2_12_12_1$ two molecules were present in the asymmetric unit (ASU), in one chain (orange) a loop encompassing residues 628-641 could not be modelled (dotted line) while in the second chain (green) this loop (purple, arrow) was ordered via interactions with the other chain. N-linked glycans are displayed as sticks on residues N544 in both chains and on N598 in the second chain only. **B.** In the $P3_221$ crystal form, four molecules of FN3-4 were arranged in a tetrameric assembly such that the loop (residues 628-641, arrows) in each chain was ordered via interactions with an adjacent chain. N-linked glycans are displayed as sticks on residues N544 and N598. **C.** Superposition of all six chains from the two crystal structures of PTPRM FN3-4.

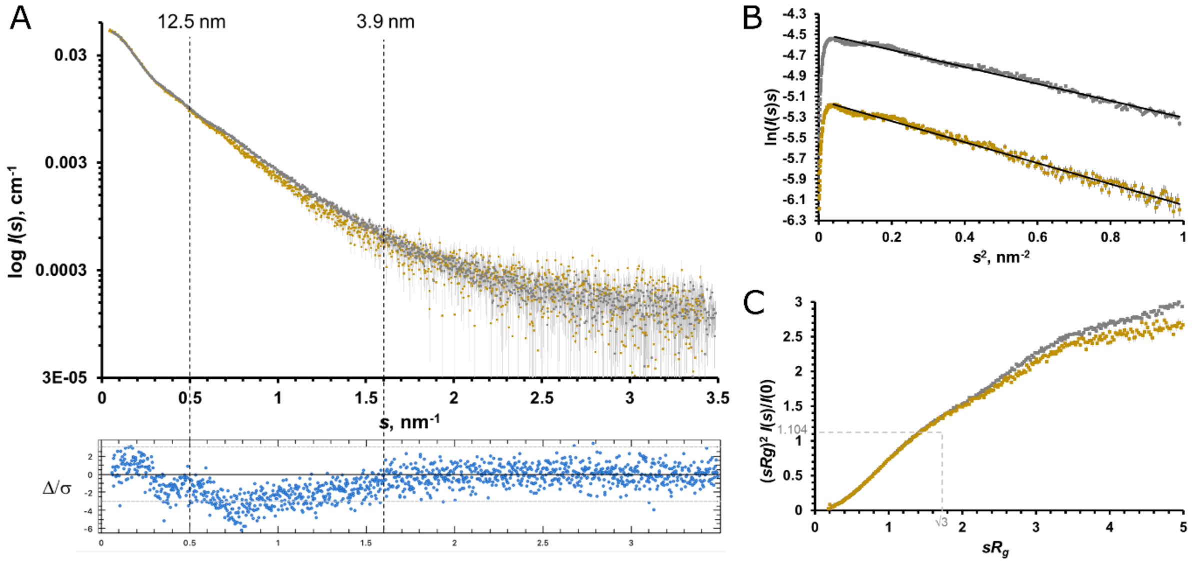


**Fig. S4. Comparison of PTPRM-ECD and PTPRK-ECD SAXS profiles. A.** Scaled SAXS data profiles measured from monomeric PTPRK-ECDs (orange) and PTPRM-ECDs (gray) showing differences in the scattering intensities at lower scattering angles, and especially between $0.5 < s < 1.6$ nm$^{-1}$ corresponding to real-space distances spanning 12.5–3.9 nm. The error-weighted residual difference plot is displayed underneath, demonstrating systematic deviations between the two datasets for $s < 1.6$ nm$^{-1}$. **B.** Modified Guinier plots used to calculate the $R_g^c$ for PTPRK-ECDs (orange) and PTPRM-ECDs (gray). The slope of the linear correlation (black line) is slightly steeper for the PTPRK-ECDs, that have a slightly larger $R_g^c$ (1.34–1.39 nm) compared to the PTPRM-ECDs (1.2–1.25 nm). **C.** Dimensionless Kratky plots calculated for PTPRK-ECDs (orange) and PTPRM-ECDs (gray) showing a monotonic increase that is typical of extended structures (unlike compact or semi-flexible modular proteins that often have a maxima in the plot near √3, 1.104).

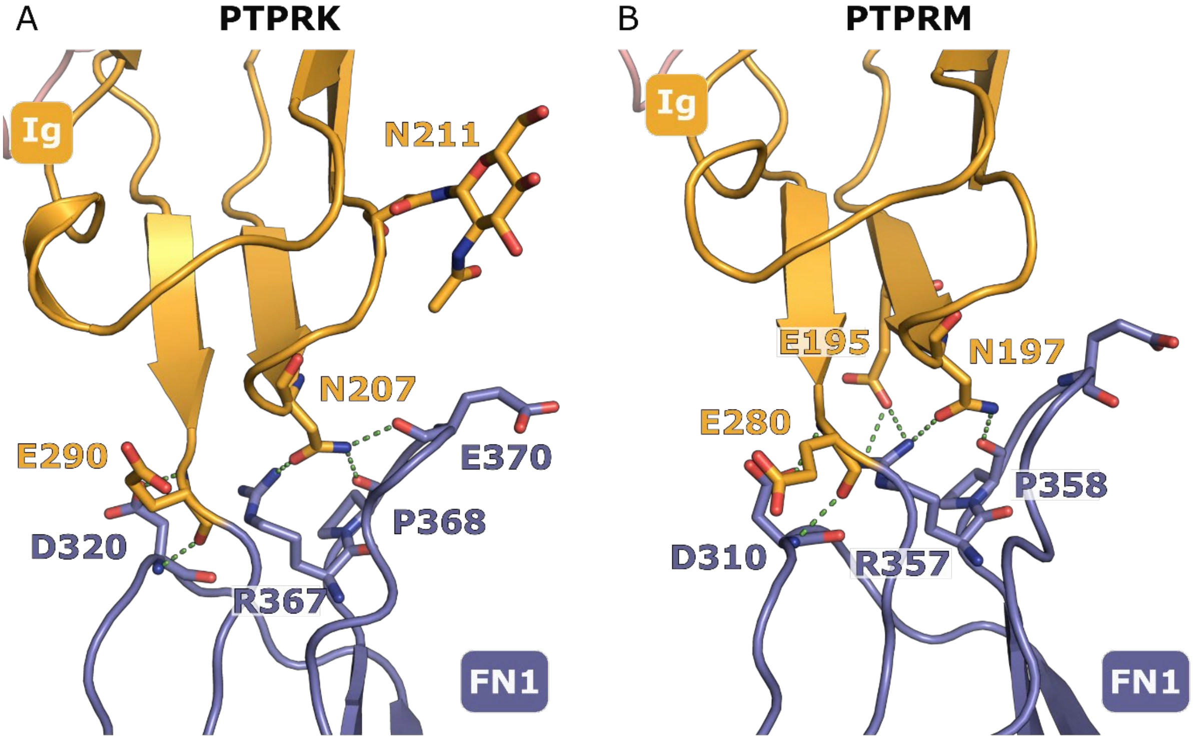


**Fig. S5. Comparison of the Ig-FN1 interface in the crystal structures of PTPRM and PTPRK. A.** Key residues involved in interdomain contacts at the interface of the PTPRK Ig (orange) and FN1 (blue) domains are shown (sticks). Hydrogen bonds between the Ig and FN1 domains are highlighted (green dashed lines). **B.** The equivalent interface in PTPRM (PDB 2V5Y) is illustrated. The numbering for PTPRM is as described in Uniprot entry P28827 and differs from the numbering in the PDB entry (by plus 20) as the deposited structure uses the post-processing numbering excluding the N-terminal signal peptide.

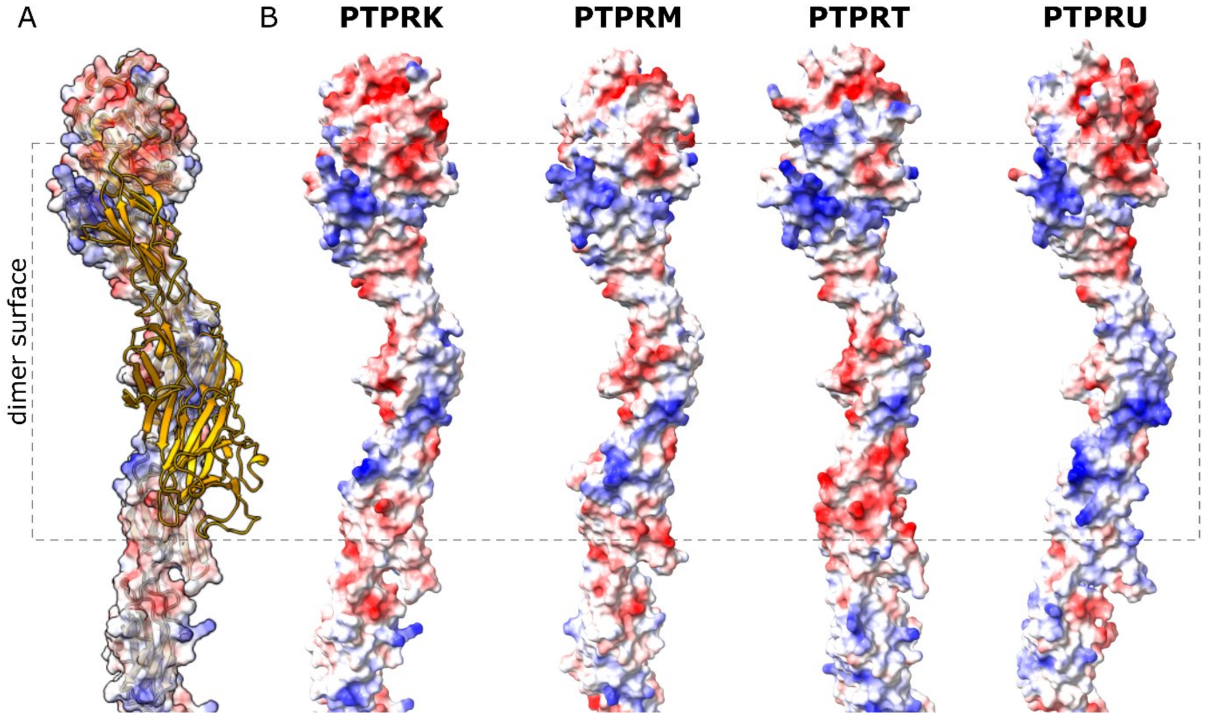


**Fig. S6. Electrostatic surface charge at the interaction interface of models for all R2B family members. A.** Surface representation of PTPRK ECD with a ribbon diagram of the second chain in the PTPRK-MAMIgFN1 dimer structure (orange) illustrating the orientation of the dimer interface. **B.** Surface electrostatics of AF2 models for the ECDs of PTPRK, PTPRM, PTPRT and PTPRU. The orientation allows direct comparison of the surface involved in dimer formation (grey dashed box).

**Table S1. X-ray diffraction data collection and structure refinement.** Data in parentheses relate to the highest resolution shell.

| | **PTPRK-MIFN1** | **PTPRM-FN3-4** | **PTPRM-FN3-4** |
|---|---|---|---|
| **Data collection** | | | |
| Beamline | I04 | I04-1 | I03 |
| Wavelength (Å) | 0.97951 | 0.9159 | 0.9763 |
| Space group | $P\,2_1\,2_1\,2_1$ | $P\,2_1\,2_1\,2_1$ | $P\,3_2\,2\,1$ |
| *Cell dimensions* | | | |
| *a*,*b*,*c* (Å) | 87.79, 93.86, 181.74 | 50.68, 89.64, 128.29 | 87.63, 87.63, 311.57 |
| α,β,γ (°) | 90, 90, 90 | 90, 90, 90 | 90, 90, 120 |
| Resolution (Å) | 79.05 – 3.00 (3.05 – 3.00)* | 128.29 – 2.89 (2.94 – 2.89) | 77.89 – 3.08 (3.14 – 3.08) |
| $R_{merge}$ | 0.079 (2.023) | 0.317 (3.531) | 0.170 (2.322) |
| $R_{pim}$ | 0.023 (0.577) | 0.093 (0.996) | 0.057 (0.819) |
| $CC_{1/2}$ | 1.000 (0.561) | 0.991 (0.432) | 0.993 (0.391) |
| $I/\sigma I$ | 18.7 (1.3) | 7.0 (0.9) | 9.1 (0.4) |
| Completeness (%) | 100 (99.1) | 100 (96.7) | 99.8 (92.0) |
| Multiplicity | 13.3 (13.1) | 12.4 (13.3) | 10.0 (8.5) |
| **Refinement** | | | |
| Resolution (Å) | 60.46 – 3.00 (3.09 – 3.00) | 73.48 – 2.89 (3.11 – 2.89) | 75.80-3.09 (3.20-3.09) |
| No. reflections | 30809 | 13636 | 26111 |
| $R_{work}/R_{free}$ | 0.228/0.274 | 0.234/0.288 | 0.243/0.290 |
| *Molecules per ASU* | 2 | 2 | 4 |
| *No. atoms* | | | |
| Protein | 5543 | 3684 | 7692 |
| Glycans | 183 | 125 | 297 |
| Other | 4 | 0 | 0 |
| *B-factors* | | | |
| Protein | 116.0 | 75.1 | 112.6 |
| Glycan | 145.7 | 94.9 | 135.8 |
| Other | 120.65 | 0 | 0 |
| *Ramachandran* | | | |
| Favoured (%) | 98.0 | 97.8 | 97.9 |
| Outliers (%) | 0.0 | 0.0 | 0.0 |
| *r.m.s. deviations* | | | |
| Bond lengths (Å) | 0.003 | 0.003 | 0.012 |
| Bond angles (°) | 0.735 | 0.779 | 1.360 |
| PDB entry | 8A1F | 8A16 | 8A17 |

**Table S2. SAXS data collection and analysis parameters**.

| **Sample details** | **PTPRK** | **PTPRM** |
|---|---|---|
| Organism | *Human* | *Human* |
| Uniprot ID (amino acid range) | Q15262 (28-752)* | P28827 (19-742) |
| SEC-SAXS buffer | 50 mM MES, pH 6.0, 250 mM NaCl, 3% v/v glycerol | |
| Sample injection volume | 40 μL | 30 μL |
| Sample injection conc. | 2.3 mg/mL | 7 mg/mL |
| SEC column | S200 Increase 5/150 | |
| SEC flow rate | 0.35 mL/min | |
| SEC temperature | 20 °C | |
| **Instrument details** | | |
| Instrument | EMBL P12 bioSAXS beam line, DESY, Hamburg | |
| Exposure time/# frames | 0.25 s (2880), entire column elution | |
| X-ray wavelength/energy | 0.124 nm (10 keV) | |
| Sample-to-detector distance | 3 m | |
| *Scattering intensity scale* | *Absolute scale, cm$^{-1}$* | |
| *SEC-SAXS primary data processing* | *CHROMIXS* | |
| # frames used for averaging | 47 | 54 |
| Working *s*-range (nm$^{-1}$) | 0.04-7.4 | 0.06-7.4 |
| **Guinier analysis** | | |
| Primary data analysis software | *PRIMUS* (*ATSAS* 3.0) | |
| Guinier *I*(0) ($\sigma$) | 0.0523(0.0001) | 0.1013(0.0002) |
| $R_g$ (Guinier, nm) ($\sigma$) | 7.0 (0.02) | 7.2(0.02) |
| $sR_g$ range/(points used) | 0.29-1.28(7-58) | 0.23-1.23(4-54) |
| $R_g^c$ (cross-section, nm) | 1.34-1.39 | 1.20-1.25 |
| ***p*(*r*) analysis** | | |
| Method | *GNOM* 5 | |
| *I*(0), POR (s) | 0.0532(0.0002) | 0.1030(0.0002) |
| $R_g$ (POR, nm) (s) | 7.5 (0.04) | 7.7(0.02) |
| $D_{max}$ (nm) | 26 | 26 |
| Quality of fit, CorMap $P$ /$\chi^2$ | 0.7/1.04 | 0.99/0.99 |
| Porod volume (nm$^3$) | 252 | 255 |
| Shape classification | extended | extended |
| **MW and hydrodynamics** | | |
| *Calculated MW, from amino acid sequence, kDa* | *81.9 (monomer)* | *81.7 (monomer)* |
| MALS protein MW, kDa | 78.5 | 80.6 |
| MALS glycan MW, kDa | 26 | 19.6 |
| MALS MW, kDa (total) | 105 | 100 |
| MW from SAXS data, kDa | 113 (106-127) | 94 (89-96) |
| **Ab initio modeling** | | |

| | | |
|---|---|---|
| Method | *DAMMIN* | |
| Symmetry | P1 | |
| #models used for averaging | 15 | 12 |
| Normalised spatial discrepancy | 0.76 | 0.77 |
| **Quality-of-fit, $\chi^2$, CorMap *P* | 1.06/0.23 | 1.02/0.65 |
| **Rigid body modeling** | | |
| Method | *AF2/X-ray/CORAL structure* | |
| Symmetry | P1 | |
| Quality-of-fit, $\chi^2$/CorMap *P* | 1.17/0.072 | 1.10/0.293 |
| Top model name | PK5su28.pdb | PM5sub6.pdb |
| **SASBDB accession codes** | SASDPF3 | SASDPG3 |

* The sequence of PTPRK used in this study is isoform 2 in the UniProt entry, possessing an alanine insertion relative to canonical sequence at position 731.

** Of final refined single model derived from the n-model cohort.
